# Phylogeny-aware linear B-cell epitope predictor detects candidate targets for specific immune responses to Monkeypox virus

**DOI:** 10.1101/2022.09.08.507179

**Authors:** Felipe Campelo, João Reis-Cunha, Jodie Ashford, Anikó Ekárt, Francisco P. Lobo

**Affiliations:** Department of Computer Science, Aston University, Birmingham, B4 7ET, UK; Department of Biology, University of York, YO10 5DD, UK; Department of Genetics, Ecology and Evolution, Universidade Federal de Minas Gerais, Belo Horizonte, Brazil

**Author notes:** These authors contributed equally to this work (Felipe Campelo, Francisco P. Lobo).

## Abstract

Monkeypox is a disease caused by the Monkeypox virus (MPXV), a double-stranded DNA virus from genus *Orthopoxvirus* under family *Poxviridae*, that has recently emerged as a global health threat after decades of local outbreaks in Central and Western Africa. Effective epidemiological control against this disease requires the development of cheaper, faster diagnostic tools to monitor its spread, including antigen and serological testing. There is, however, little available information about MPXV epitopes, particularly those that would be effective in discriminating between MPXV infections and those by other virus from the same family. We used the available data from the Immune Epitope Database (IEDB) to generate and validate a predictive model optimised for detecting linear B-cell epitopes (LBCEs) from *Orthopoxvirus*, based on a phylogeny-aware data selection strategy. By coupling this predictive approach with conservation and similarity analyses, we identified nine specific peptides from MPXV that are likely to represent distinctive LBCEs for the diagnostic of Monkeypox infections, including the independent detection of a known epitope experimentally characterised as a potential specific diagnostic target for MPXV. The results obtained indicate ability of the proposed pipeline to uncover promising targets for the development of cheaper, more specific diagnostic tests for this emerging viral disease. A full reproducibility package (including code, data, and outputs) is available at https://doi.org/10.5281/zenodo.7838331.

## 1 Introduction

Monkeypox is a zoonotic viral disease caused by the Monkeypox Virus (MPXV), a pathogen endemic to Western and Central Africa and a member of the genus *Orthopoxvirus* (OPXV), a clade that also hosts the Variola (Smallpox) virus. MPXV was initially found to infect humans in the Democratic Republic of Congo in 1970 [1,2]. Since then, two lineages have been identified: the Central Africa clade (more lethal, with mortality of 8.7%, usually referred as clade 1) and the Western Africa clade (less lethal, with mortality of 3.6%, nowadays referred as clades 2 and 3) [3]. Outbreaks of all lineages have been reported in Africa, Asia, North America and Europe [4].

The ongoing MPXV outbreak started in early May 2022 and is the largest one described thus far in non-endemic countries, comprising more than 52,000 officially recorded cases as of September 2nd 2022, 99% of them in countries that have not historically reported Monkeypox [5]. The current epidemic has been identified in 100 countries across the five WHO regions. The circulating lineage is a member of clade 3 MPXV, and is most closely related to a large outbreak occurring in Nigeria in 2017–2018 [6].

Smallpox vaccination throughout the twentieth century was routinely done with the Vaccinia virus, an *Orthopoxvirus* closely related to Smallpox, which also provided approximately 85% of protection against MPXV [7]. Even though this vaccine provides lingering protective immunity that can last for decades [8], the end of routine vaccination against Smallpox after the eradication of the disease in 1980 means that it has now been more than forty years since widespread vaccination promoted Smallpox (and, by extension, Monkeypox) immunity [9], which may partly explain the recent surge in Monkeypox cases across the globe.

Experience with large-scale testing has been acquired during the ongoing Coronavirus Disease 2019 (Covid-19) pandemic, with national health services now providing standard testing protocols. Another lesson learned during this pandemic is the importance of cheaper and faster testing alternatives: direct survey for viral antigens (AGs) as an indication of active infection through several simple testing kits has greatly improved the control of the Covid-19 pandemic [10]. In the case of MPXV, to the best of our knowledge, all such current testing protocols rely on nucleic acid amplification through polymerase chain reaction (PCR) [11,12], a procedure that is considerably more time-consuming and harder to deploy at scale than AG-based tests [13]. Even though the development of tests based on the presence of antibodies (ABs) specific for MPXV AGs does not provide useful information for the detection of active infections, these tests are of utmost importance for epidemiological studies, as has been the case in the ongoing Covid-19 pandemic [14]. In the ongoing MPXV outbreak, specific AB-based testing would be useful to distinguish previous MPXV infections from the ABs resulting from, e.g., smallpox vaccination or earlier infections from other related viruses.

The computational prediction of linear B-cell epitopes (from now on referred as LBCEs), a scientific field under active development for more than 40 years [15], has become an important step in the development of vaccines and diagnostic tests for infectious diseases [16–18]. A common premise in this field is the development of generalist predictors, defined as models trained with a taxonomically diverse set of entries and intended to generalise to any epitope prediction task, irrespective of the pathogen being studied. Recently, our group demonstrated how organism-specific training generates consistent improvements over generalist methods for LBCE prediction [19].

The Immune Epitope Database (IEDB) [20] is arguably the most complete repository of curated epitopes^1^ and non-epitopes and associated metadata, and spanning thousands of source species including viruses, protozoans and other classes of parasites. A considerable fraction of this data comprises LBCEs, which are the main focus of this study. Currently, IEDB provides very limited data describing LBCEs associated with MPXV, with only five peptides representing epitope-containing regions described thus far. Such knowledge is of uttermost importance for the development of AG- and AB-based tests, and to aid in the rational design of sub-unit, peptide and chimeric vaccines. The scarcity of epitope information for MPXV contrasts with the relatively larger availability of information about B-cell epitopes for the taxonomic lineage where MPXV is found: after initial processing, IEDB was found to provide 20, 66 and 128 peptides labelled as epitopes in genus *Orthopoxvirus*, class *Pokkesviricetes* and kingdom *Bamfordvirae*, respectively, with a number of labelled counter-examples also available (14, 18 and 39, respectively).

In this work we deploy a phylogeny-aware modelling approach to detect LBCE-containing regions in MPXV proteins, using IEDB data filtered at distinct taxonomic levels to train predictive models for the detection of LBCEs in *Orthopoxvirus* proteins. We demonstrate that the selected model, trained on data from all viruses under the *Bamfordvirae* kingdom and optimised for prediction of *Orthopoxvirus* epitopes, substantially outperforms the current standard LBCE prediction tool, Bepipred 2.0 [21] with a similarly optimised threshold.

The selected model is applied to predict LBCE on 190 MPXV protein sequences from a viral lineage of the current outbreak. The resulting predictions are evaluated for conservation across MPXV genomes publicly available, and also for dissimilarity from the predicted proteomes of humans, other *Poxviridae*, and other pathogens that may result in clinical presentations similar to Monkeypox infections. This analysis yielded nine peptides which are highly conserved in MPXV and also dissimilar from sequences from the host and other pathogens. These peptides represent promising targets for the development of AG or serological diagnostic tests of Monkeypox.

## 2 Methodology

### 2.1 Data Sets

Figure 1(A) illustrates the process of retrieving and processing the relevant data sets.

**Fig 1.**
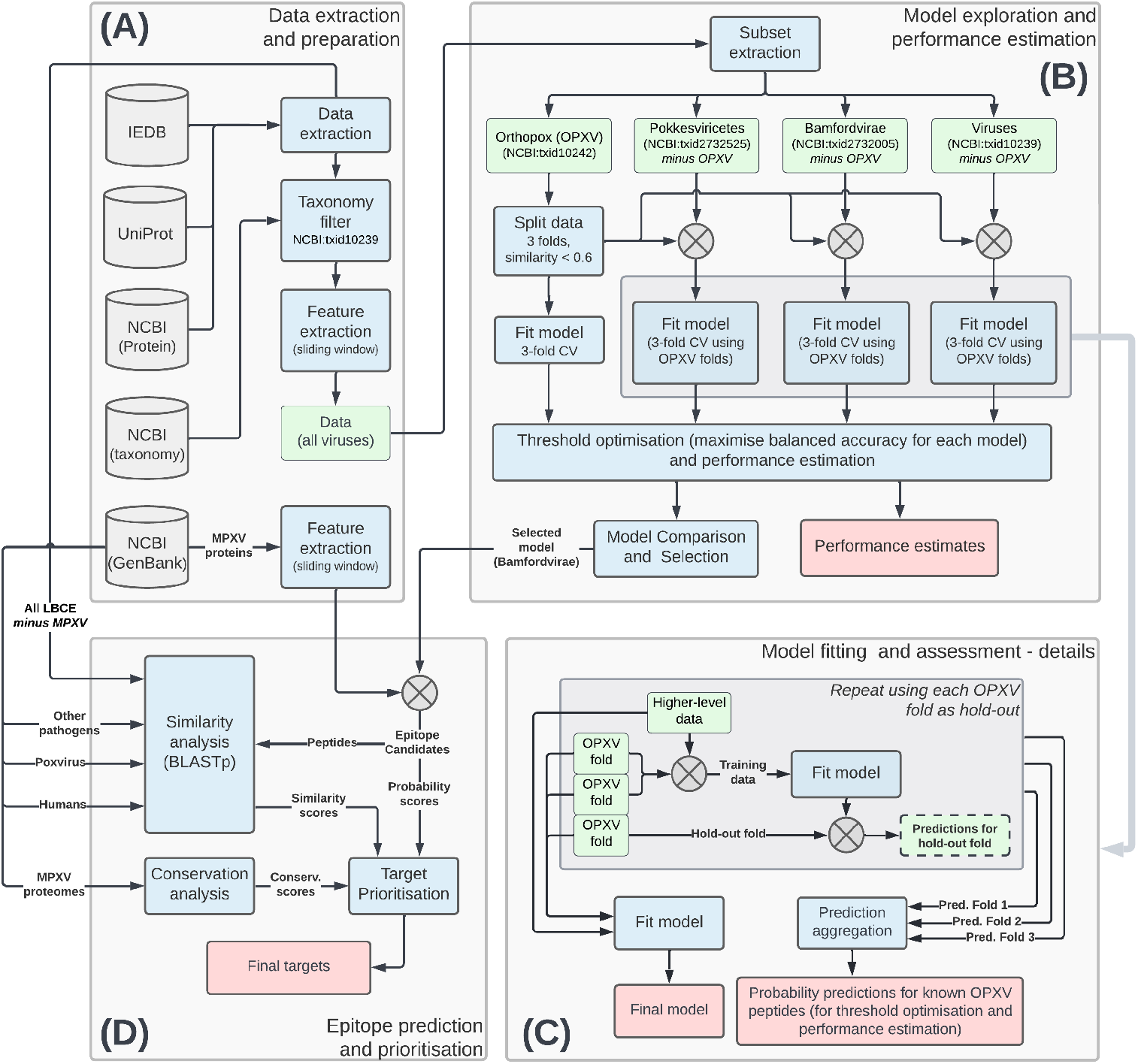
Experimental protocol for the development and selection of OPXV-specific models and the prediction and prioritisation of candidate epitopes for MPXV. (A) Data sources and dataset generation process. Queries to the IEDB, UniProt and NCBI databases were automated using the development version of R package *epitopes* [22]. The MPXV proteins were retrieved from the Genbank file corresponding to the first complete genome sequence of the current MPXV outbreak (isolate MPXV USA 2022 MA001, accession number ON563414). (B) Model development and assessment. The detailed view of the model training process (C) highlights the use of OPXV folds for model training and estimation of model performance. (D) Use of the selected model (re-trained with all OPXV examples after the performance assessment, as shown in block C) to predict epitopes on MPXV proteins. The candidate peptides are ranked based on predicted probability, and a similarity profile is estimated from local alignment against all available epitopes from superkingdom *Viruses*.

#### 2.1.1 Training data

Data extraction, filtering and consolidation was performed using functions available in the development version of R package *epitopes* [22], based on the full XML export of the IEDB database [20] on 20 May 2022. All entries identified as LBCEs from organisms under the superkingdom *Viruses* (NCBI:txid10239) were extracted from the IEDB export, with associated proteins retrieved from the NCBI protein database [23] and Uniprot [24]. Peptides were labelled as positive if half or more of the assays associated with that IEDB entry reported a positive result, and positive-labelled peptides of length greater than 30 amino acid residues were removed, to prevent long “Epitope-containing regions” from adding excessive noise to the training data. Overlapping peptides of the same class were merged into a single entry to prevent partial data duplication. The resulting sequence data was then tabularised using a sliding window strategy (window length = 15, step size = 1) [19], and a set of 385 statistical and physio-chemical features based on the local neighbourhood of each residue were calculated.^2^

The resulting data set containing information on all labelled peptides related to viral LBCEs was further divided as follows: All entries related to pathogens under genus *Orthopoxvirus* (OPXV) (NCBI:txid10242) were isolated in a separate data set, and a similarity-aware data splitting strategy was used to split this data into three folds based on normalised local alignment scores.^3^ Four of the positive examples available under the OPXV data correspond to known MPXV epitopes (IEDB epitope IDs 10135, 7309, 39258 and 73976), with no recorded negative MPXV examples.

The remaining data *(excluding OPXV*) were used to generate three data sets, with all entries related to pathogens (i) under class *Pokkesviricetes* (NCBI:txid2732525); (ii) under kingdom *Bamfordvirae* (NCBI:txid2732005); and (iii) under the superkingdom Viruses (NCBI:txid10239).^4^ Table 1 summarises the main characteristics of these data sets.

**Table 1.** Data sets used for development and validation of the Orthopoxvirus-specific LBCE predictor.

| Dataset | Labelled peptides (-/+) | Number of proteins | Details |
| --- | --- | --- | --- |
| *Orthopoxvirus* (OPXV)<br>*NCBI:txid10242* | 14- / 20+ | 15 | Split in three folds based on normalised protein local alignment scores. Smith-Waterman normalised similarity threshold: 0.6 |
| *Pokkesviricetes*<br>*NCBI:txid2732525* | 18- / 66+ | 36 | Excluding all OPXV entries. |
| *Bamfordvirae*<br>*NCBI:txid2732005* | 39- / 128+ | 71 | Excluding all OPXV entries. |
| *Viruses*<br>*NCBI:txid10239* | 10,167- / 6307+ | 5493 | Excluding all OPXV entries. |

#### 2.1.2 Monkeypox proteins

Existing annotation data related to MPXV proteins was retrieved from the Genbank file ON563414 (field “note”). In total, 190 proteins were retrieved and processed similarly to the training data, as described in section 2.1.1 and Fig. 1(A).

### 2.2 Modelling

Fig. 1(B) and (C) detail the training and assessment of models for LBCE prediction in pathogens under genus *Orthopoxvirus*. As established by Ashford et al. [19], pathogen-specific training has been shown to improve the performance of LBCE predictors when compared to generalist models trained with larger, but substantially more taxonomically heterogeneous, data. As the number of available MPXV-specific examples in IEDB is very low to train, or even properly assess, any predictive models,^5^ we used the same rationale as Ashford et al. [19] to investigate models that generalise well to our genus of interest, i.e., models displaying good predictive ability to detect *Orthopoxvirus* LBCE.

Random Forest (RF) models were trained using the datasets described in Subsection 2.1.1. These datasets present increasing amounts of data, obtained by expanding the data inclusion criterion to admit viral lineages with increasing taxonomic distance from OPXV. The model trained using only the OPXV data was assessed using 3-fold cross-validation with similarity-based folds as described in Subsection 2.1.1. The models trained with data from higher taxonomic levels were also assessed using 3-fold cross-validation, with each sub-model trained on the combination of the full higher-level data set plus two folds of the OPXV data, and deployed to predict the examples in the third OPXV fold. These predictions were then aggregated together for threshold optimisation and performance comparison. After the predicted probabilities (residue-wise) for each hold-out fold were aggregated and recorded, a final model was then generated for each of the datasets consisting of the full OPXV data combined with one set of higher-level observations *(Pokkeviricetes, Bamfordvirae*, and *Viruses*, respectively). This process is detailed in Fig. 1(C).

The vectors of predicted probabilities for each residue of the OPXV data were then used to optimise the classification threshold for each model. This process was done by simple exhaustive search, varying the threshold from 0.01 to 0.99 in increments of 0.01, and calculating the balanced accuracy (simple mean of model specificity and sensitivity) at each level. The threshold values that resulted in the highest balanced accuracy were selected for each model.

All RF models were trained using the implementation from R package *ranger* [26] version 0.13.1 with standard hyperparameters. Class imbalance was treated by stratified undersampling of the majority class, ensuring that all peptides remained represented by at least one observation in the re-balanced data.

The trained models are applied to predict the LBCE probability for each residue of a given protein, and return a probability value for each protein position. Contiguous sequences of at least eight residues above the optimised model threshold were considered as potential epitope-containing peptides, with the probability score of the peptide attributed as the mean of the predicted epitope probabilities of its individual amino acid residues.

### 2.3 Conservation and Similarity Evaluation

The conservation of the candidate peptides in the predicted proteome of 88 Monkeypox isolates (Supplementary file 5) was estimated using BLASTp [27] with options `-seg no -evalue 10000 -word_size 3` (optimized for short sequences). Similarly, the dissimilarity between each of the predicted peptides and the predicted proteomes of:

- *Homo sapiens* (GCF_000001405.40);
- *Sarcoptes scabiei* (GCA_000828355.1), Measles morbillivirus (GCF_000854845.1) and *Treponema pallidum* (GCF_000246755.1), pathogens with potentially similar clinical presentations [1]; and
- All sequences available from NCBI under family *Poxviridae* and not identified as Monkeypox (a total of 576 isolates of 72 species)

was estimated using a normalised dissimilarity index yielding values in the [0,1] range, *diss* = 1 — *(identity × coverage/*10^4^). All resulting BLASTp files and identifiers of all sequences are provided in the Supplementary file 6.

## 3 Results

### 3.1 Phylogeny-aware modelling yields superior predictive performance for OPXV

The performance obtained by the models trained on each data set for the prediction of OPXV epitopes as well as the performance observed for Bepipred 2.0 [21] was quantified using eight commonly used performance indicators: area under the ROC curve (AUC), F1 indicator (F1), Matthews Correlation Coefficient (MCC), Balanced accuracy, Sensitivity and Specificity, and the Positive (PPV) and Negative (NPV) predictive values [28]. The first four (AUC, F1, MCC and Balanced accuracy) represent aggregated measures of performance, whereas the remaining ones (Sensitivity, Specificity, PPV and NPV) quantify specific predictive properties. Standard errors for each value, as well as p-values for individual comparisons against the Bepipred 2.0 baseline, were estimated using bootstrap.^6^

Several interesting observations emerge from the results illustrated in Fig. 2. First, it is clear that performance increases across most performance metrics as more data from phylogenetically-related pathogens is added to the training set, but stabilises or even decreases once the much larger volume of data from unrelated viruses is added into the training set. This result reinforces and expands some observations discussed in our earlier work [19], particularly the idea that training bespoke models based on data more directly related to the pathogen (or, in the particular case of this experiment, genus) under study improves the model’s predictive ability. Moreover, it is clear that the performance of the bespoke models is generally superior to that of Bepipred 2.0, a generalist predictor commonly used as the standard computational tool for LBCE prediction. The main exception to this observation is the model trained exclusively with the OPXV data (or, more precisely, with 2/3 of the available OPXV data on IEDB). Although this underperformance may seem superficially at odds with the notion of organism-specific model training, we suggest that the extremely low number of OPXV examples may be insufficient to fit models with good predictive abilities. This finding reinforces results obtained in our preliminary exploration of the limits of organism-specific training: at the very low end of data availability the resulting models may not achieve sufficient generalisation ability [31]. It is also interesting to note that the bespoke models (with the exception of the one trained on all viral epitopes) present lower specificity than Bepipred 2.0, but non-inferior PPV and considerably higher sensitivity and NPV. This indicates that the modelling approach employed here led to models that are better capable of correctly discarding non-immunogenic peptides (NPV) without sacrificing their precision in detecting epitopes (PPV), when compared to Bepipred 2.0.

**Fig 2.**
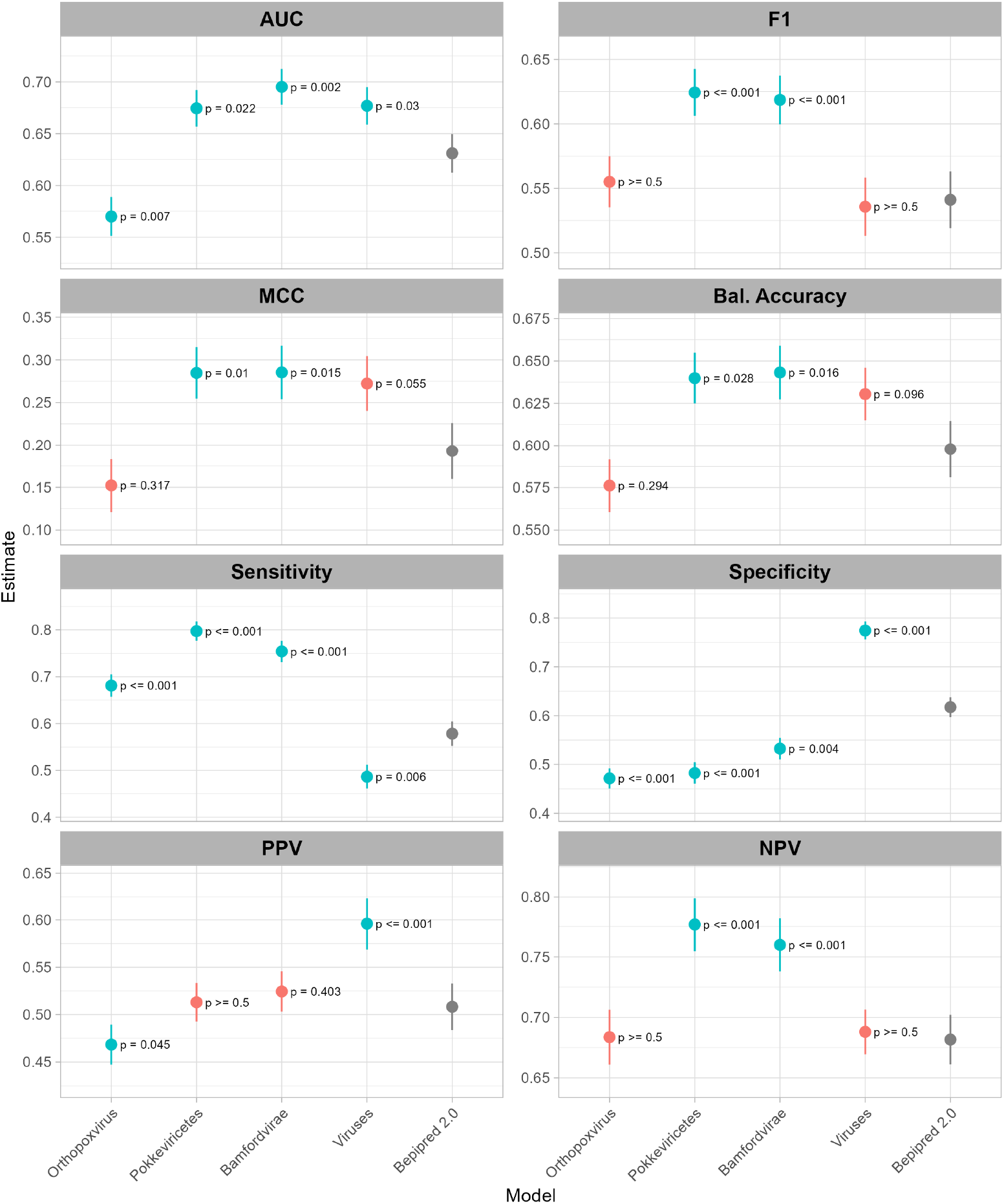
Performance assessment of all models using the OPVX peptides currently available in IEDB as the test set (see Fig. 1 B-C for details). The performance attained by Bepipred 2.0 is added as a comparison baseline. All models, including Bepipred 2.0, used thresholds optimised to maximise balanced accuracy on all labelled OPXV peptides available from IEDB. Vertical bars represent bootstrapped estimates of the standard errors. The p-values reported in the panels refer to performance comparisons of each model against the Bepipred 2.0 baseline, also estimated using bootstrap resampling. Statistical significance at the 95% confidence level is indicated in green.

Based on the results illustrated in Fig. 2, the model trained on data from kingdom *Bamfordvirae* was selected as the one most likely to provide good predictive performance for MPXV epitopes, given that it provided the highest values of all aggregated performance indicators (AUC, F1, MCC and Balanced Accuracy), systematically outperforming Bepipred 2.0. The optimised probability threshold for this model was determined as 0.54. The performance level observed for this model is expected to lead to more reliable predictions in terms of precision in the determination of epitopes and, critically for pathogens with relatively small genomes such as MPXV, without providing overly conservative predictions that could discard valuable targets for downstream experimental characterisation aiming at developing diagnostic tests and vaccines.

### 3.2 Selected model predicts new LBCE candidates for MPXV

The selected model (trained on all available examples under kingdom *Bamfordvirae* and optimised to maximise balanced accuracy on *Orthopoxvirus* data) was applied to generate predictions for the 190 proteins derived from the Genbank file corresponding to the first complete genome sequence of the current MPXV outbreak (isolate MPXV_USA_2022_MA001, accession number ON563414).^7^

The selected model predicted 1,241 unique peptides with probabilities above the optimised model threshold. Supplementary file 2 provides a summary of the number of peptides predicted as epitope-containing regions for each protein, as well as the maximum peptide probability found in each protein. Supplementary file 4 provides further details of 66 unique peptides selected for further analysis, as detailed in Subsection 3.3.

### 3.3 Conservation and similarity analyses suggest potential epitopes for diagnostic tests

From the predictions provided by the selected model and on the similarity to known non-MPXV LBCE extracted from the IEDB, 66 unique peptides presenting a predicted probability greater than 0.75 and similarity score to known epitopes lower than 0.75 were selected for further analysis, aimed at identifying peptides representing interesting targets for the development of diagnostic tests with reduced probability of cross-reactivity against other pathogens. To that end, sequence conservation of candidates among MPXV isolates as well as dissimilarity to proteins from humans, other *Poxviridae* and selected pathogens with similar clinical presentations to Monkeypox were assessed. Epitope conservation was evaluated in all 88 Monkeypox isolates with annotated predicted proteomes available in NCBI on 24 August 2022, whereas dissimilarity to the host and other pathogens was assessed using the data sources described in Subsection 2.3.

Figure 3 illustrates the results of this analysis. Of the sixty-six unique peptides pre-selected based on the probability score attributed by the model and their similarity score to known LBCE extracted from the IEDB (Fig 3, top), nine displayed high dissimilarity to human proteins (≥ 0.5) and a dissimilarity to Poxvirus greater than a minimal threshold of 0.05 (Fig. 3, bottom),^8^ as well as reasonably high conservation scores (≥ 0.75).

**Fig 3.**
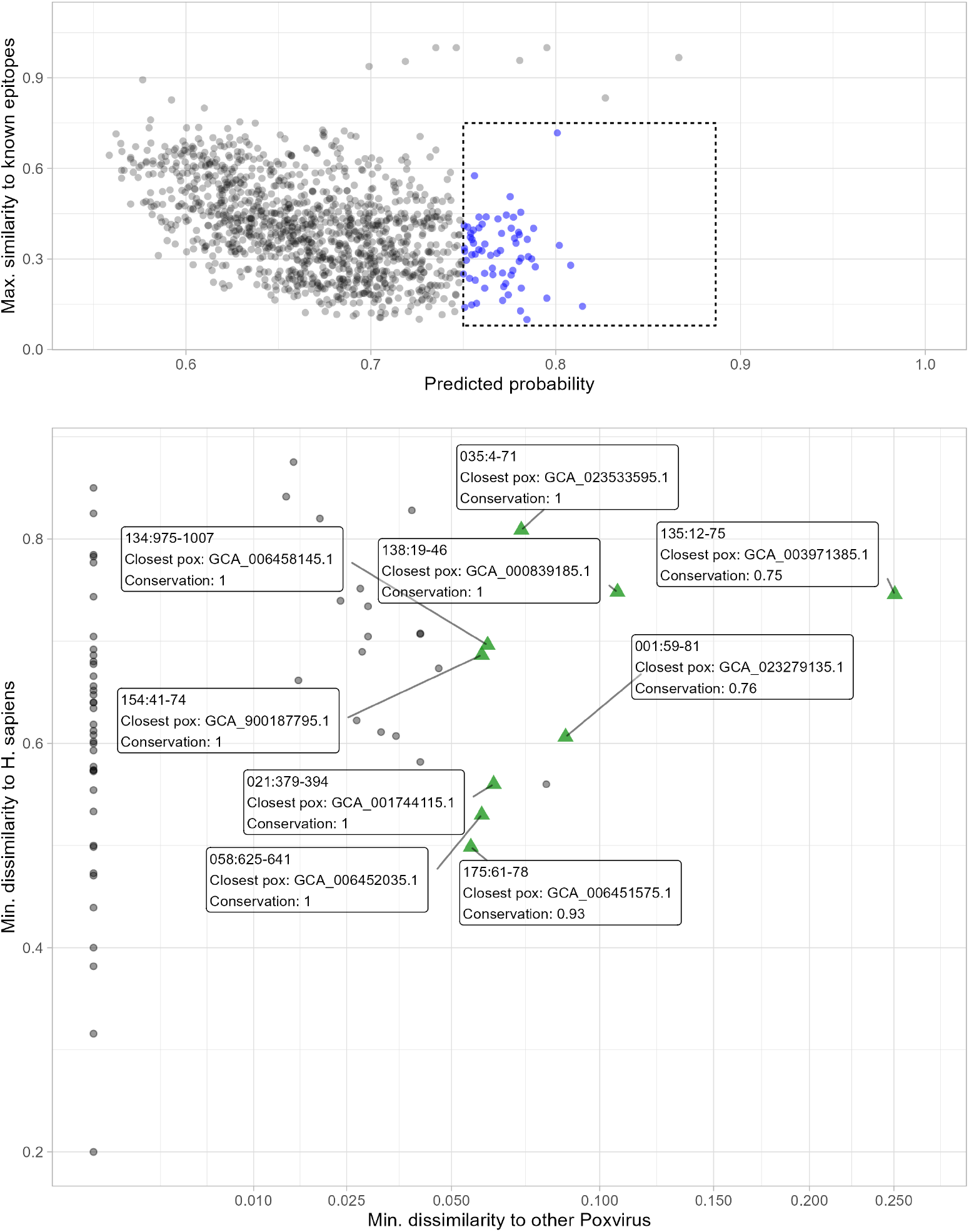
**Top**: Predicted peptides in the space of *predicted probability × max. similarity to known LBCE*. Predictions with low similarity and high probability were highlighted and selected for further analysis. Highlighted points were projected (**Bottom**) onto the space of Min. dissimilarity to proteins from *other Poxvirus × H. sapiens*. Candidates with perfect or near-perfect similarity with other Poxvirus (left side) would be unsuitable for the development of diagnostic tools to differentiate MPXV infections from those pathogens. Predicted LBCE with a conservation score ≥ 0.75, dissimilarity to human proteins ≥ 0.5 and dissimilarity to other Poxvirus proteins ≥ 0.05 are highlighted as green triangles as candidates for diagnostic test development.

We propose that these nine peptides, highlighted as green triangles in the figure, may represent interesting targets for the development of diagnostic tests with good ability to differentiate between MPXV and other infections. These peptides are found in proteins fulfilling a variety of roles in MPXV biology and encompassing both early and post-replicative expressed genes. These roles range from virulence factors secreted by infected cells that inhibit molecular hubs of the host immune response to structural components of viral particles and enzymes playing roles in viral DNA replication (Supplementary file 8). Importantly, one of the candidate epitopes (identified in Table 2 as 138:19-46) coincides with a peptide previously identified as a highly specific MPXV epitope [32] (`teFFSTKAAKNPETKREAIVKAYGDDNEETlkq`, with uppercase characters highlighting our prediction). Detection of this epitope, which was not part of the training set since it is not present in IEDB, corroborates both the ability of the selected model to accurately detect LBCE and the usefulness of the predictive pipeline developed in this work in identifying valuable targets for the development of diagnostic tests. Table 2 provides further details of all predicted LBCE identified as potential diagnostic targets.

**Table 2.**
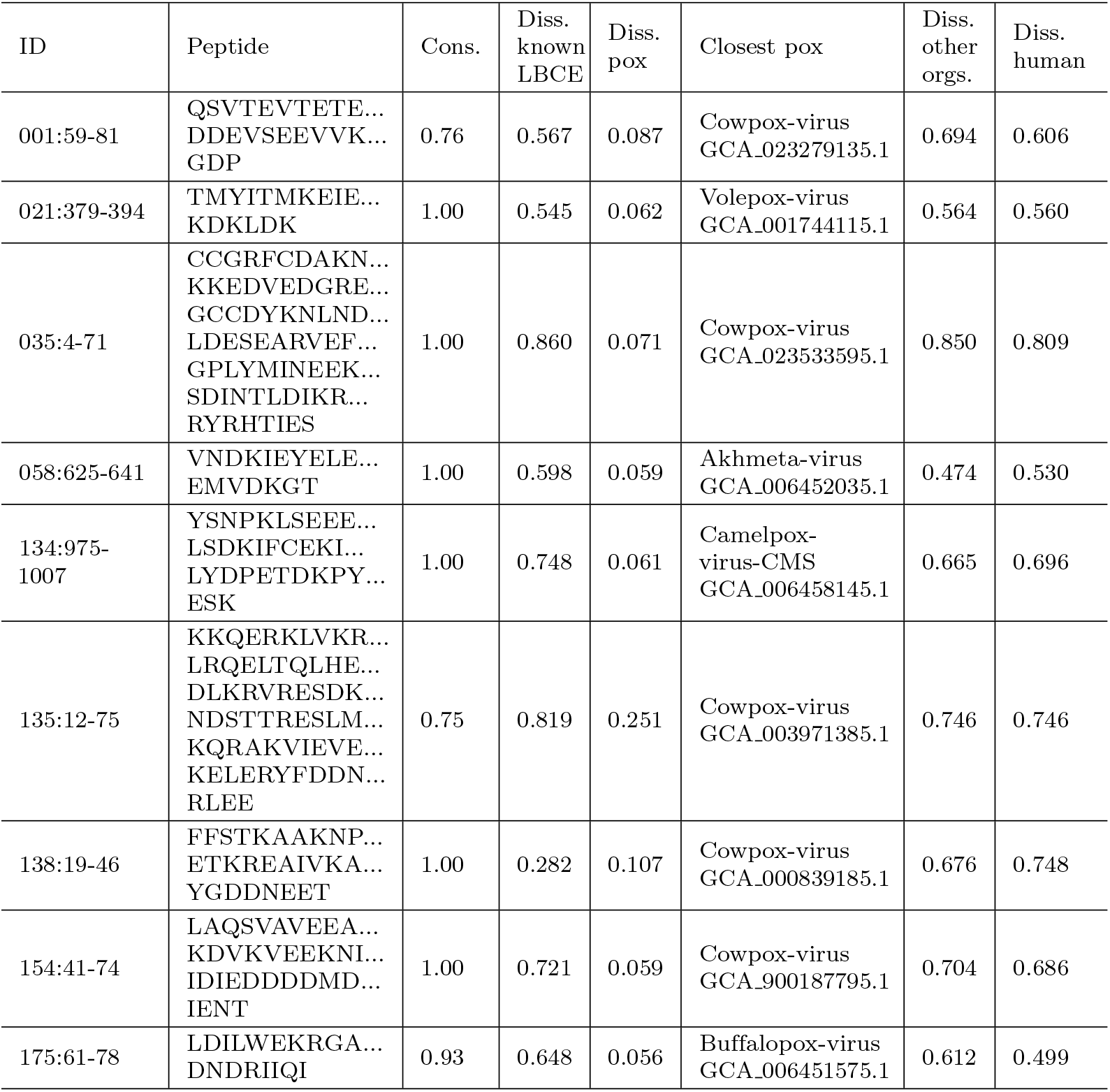
Selected peptides as potential LBCE-containing regions for diagnostic test development. ID indicates the protein number ID and position (see supplementary files 2 and 3 for details). Candidate 138:19-46 is known to be an epitope with potential diagnostic applications [32], and was discovered independently in this work, highlighting the ability of the proposed pipeline to uncover MPXV LBCEs. Note: *Diss. other orgs*. refers to the minimum dissimilarity score to the non-Poxvirus pathogens evaluated.

## 4 Discussion

By combining a specific LBCE predictor built under a phylogeny-aware framework with sequence conservation among known MPXV isolates and the dissimilarity of predicted targets to sequences from humans, other Poxvirus and infections with similar clinical symptoms, we were able to select nine peptides as potential candidates for the AG- and AB-based diagnostic of current or previous MPXV infections, respectively. The dissimilarity between candidate peptides and proteins from humans and from non-pox pathogens was higher than that observed for other *Poxviridae* family members (Table 2). This result is expected due to the higher sequence conservation of *Orthopoxvirus* as well as other viruses under the *Poxviridae* family, as a consequence of a more recent common ancestor. In practical terms, this could represent a limitation for specie-specific AG- and AB-based diagnostic tests of MPXV. On the other hand, the peptide from protein A29L (which appears in the present study under the ID *MPXV-USA_2022_MA001-138.t01*) have been shown to be a promising target for the specific diagnostic of MPXV, even with variations in only a few key residues [32]. It is important to highlight that the peptide identified by Hughes et al. [32] also appears in the list of potential diagnostic LBCEs identified by our pipeline, highlighting the ability of the proposed method to effectively identify LBCEs from MPXV protein sequence data.

Beyond the detection of potential Monkepox LBCEs, the phylogeny-aware approach used in this work has been shown to provide better results than the current standard tool for LBCE prediction (Bepipred 2.0), reinforcing earlier results that already suggested this to be the case [19]. More importantly, the successful development of model tailored specifically for an emerging global pathogen with almost no LBCE data available shows that this approach is useful even beyond the lower bounds investigated in an earlier work [31], and that utilising information from phylogenetically-related pathogens can be a successful strategy for the development of predictive models tailored for specific groups of pathogens that share a common ancestor. With the current wide availability of considerable computing power even on low-cost and mobile devices, there is clearly scope for a wider use of pathogen-specific models, trained and tuned for the prediction task of interest.

It is important to highlight that, even though not surveyed in this work, a conceptually similar search strategy could be used to help selecting immunogenic peptides for rational vaccine design. This can be achieved, for instance, by modifying the prioritisation strategy to focus on peptides with high conservation scores across all *Orthopoxvirus* (including recent MPXV variants) or even across all *Poxviridae* of medical interest, as a strategy to investigate potential pan-vaccines against multiple pathogens. As long-term protective B-cell immune response is observed in smallpox vaccination in humans [33,34] and the antibody immune response is known to be crucial in controlling Poxvirus secondary infections in mice [35], it is likely that a pan-vaccine based on highly conserved, common epitopes across multiple *Poxviridae* - which could be detected by the proposed pipeline with minimal modifications - could provide immunity that is not only cross-protective across multiple health threats, but also potentially less likely to lose efficacy from continuous virus evolution.

In conclusion, the use of a phylogeny-aware modelling approach trained on available LBCE data for MPXV and other phylogenetically-related pathogens, in conjunction with conservation and similarity analyses of the candidate epitopes detected, points to specific peptides with a high likelihood of eliciting an MPXV-specific immune response in humans. The fact that the pipeline was able to independently detect a known, experimentally confirmed epitope with the desired characteristics that was not part of the model’s training set [32] is important, as it provides external validation of the methodology used. The potential of differentiating MPXV immunity from that elicited by present or past infections from closely-related pathogens and previous smallpox vaccination is of great importance for the control and epidemiological understanding of the ongoing Monkeypox outbreak.

## Acknowledgments

Computational experiments were run using Aston University’s EPS Machine Learning Server, funded by the Engineering and Physical Sciences Research Council (EPSRC), UK. J.A. was supported by the EPSRC (Ph.D. fees and stipend). J.R.-C. was supported by the Medical Research Council under MRC New Investigator Research Grant MR/T016019/1.

## Footnotes

1 In this work we use the term *epitope* to refer both to exact LBCE and to LBCE-containing regions, as annotated in the IEDB metadata.

2 See *Supplementary File 1* for full list of features used.

3 Smith-Waterman optimal alignment [25] was used (with the BLOSUM62 substitution matrix) since the number and length of sequences to be aligned was quite small. Splits with similar characteristics could be obtained using, e.g., BLAST.

4 Other intermediate taxonomic levels - e.g., family *Poxviridae* (NCBI:txidlO24O) or order *Chitovirales* (NCBI:txid2732527) - resulted in no addition of new data from IEDB, and were therefore not considered.

5 Five peptides representing epitope-containing regions of MPXV, of which only four meet our inclusion criteria detailed in section 2.1.1; no negative examples of MPXV available.

6 The bootstrap methodology was validated by comparing the standard errors and p-values obtained for AUC against a non-resampling, non-parametric alternative [29] implemented in R package *pROC* [30].

7 See *Supplementary file 2* for details on the proteins and Supplementary *file 3* for the sequences in FASTA format.

8 The dissimilarity score of the selected peptides to the non-Poxvirus pathogens was, as expected, considerably higher than the similarity to other Poxvirus. In all cases the score was greater than 0.4.

## Notes

https://doi.org/10.5281/zenodo.7838331

### Competing Interest Statement

The authors have declared no competing interest.

### Summary of Updates

Minor corrections to links contained in the manuscript.

