## Supplementary material for "Phylogeny-aware linear B-cell epitope predictor detects candidate targets for specific immune responses to Monkeypox virus": Supplementary 1 - feature list.pdf

### Supplementary File 1

Full list of features calculated

September 8, 2022

All features were calculated based on a sliding window projection of the data, which represents each position by a 15-position wide window centred at each residue (truncated at both ends of the protein).

Feature calculation was performed using function *calc_features()* from R package *epitopes* [1], which draws heavily from implementations available in R package *protr* [2]. See the documentation of *calc_features()* for further details.

#### List of features

- **Entropy**: Sequence entropy (1 feature).
- **MolWeight**: Sequence molecular weight (1 feature).
- **AAtypes**: Proportion of amino acids of each of 9 types: tiny, small, acidic, basic, aliphatic, aromatic, polar, non-polar, charged (9 features).
- **Atoms**: Number of atoms of C, N, H, O, S in the sequence (5 features).
- **AAC**: Proportion of each natural amino acid in the sequence (20 features).
- **CTD**: Composition-Transition-Distribution (CDT) features [2] (147 features).
- **BLOSUM**: BLOSUM-derived descriptors with substitution matrix BLOSUM62, k = 5, lag = 3 and scale = TRUE [2] (75 features).
- **SOCN**: Sequence-order-coupling number with maximum lag = 3 [2] (6 features).
- **QSO**: Quasi-sequence-order descriptors with maximum lag = 3 and weighting factor w = 0.1 [2] (46 features).
- **ScalesGap**: Scales-based descriptors derived from Principal Components Analysis with pc = 5 and lag = 3 [2] (75 features).
